# Nature over Nurture: Functional neuronal circuits emerge in the absence of developmental activity

**DOI:** 10.1101/2022.10.24.513526

**Authors:** Dániel L. Barabási, Gregor F. P. Schuhknecht, Florian Engert

## Abstract

During development, the complex neuronal circuitry of the brain arises from limited information contained in the genome. After the genetic code instructs the birth of neurons, the emergence of brain regions, and the formation of axon tracts, it is believed that neuronal activity plays a critical role in shaping circuits for behavior. Current AI technologies are modeled after the same principle: connections in an initial weight matrix are pruned and strengthened by activity-dependent signals until the network can sufficiently generalize a set of inputs into outputs. Here, we challenge these learning-dominated assumptions by quantifying the contribution of neuronal activity to the development of visually guided swimming behavior in larval zebrafish. Intriguingly, dark-rearing zebrafish revealed that visual experience has no effect on the emergence of the optomotor response (OMR). We then raised animals under conditions where neuronal activity was pharmacologically silenced from organogenesis onward using the sodium-channel blocker tricaine. Strikingly, after washout of the anesthetic, animals performed swim bouts and responded to visual stimuli with 75% accuracy in the OMR paradigm. After shorter periods of silenced activity OMR performance stayed above 90% accuracy, calling into question the importance and impact of classical critical periods for visual development. Detailed quantification of the emergence of functional circuit properties by brain-wide imaging experiments confirmed that neuronal circuits came ‘online’ fully tuned and without the requirement for activity-dependent plasticity. Thus, we find that complex sensory guided behaviors can be wired up by activity-independent developmental mechanisms.

## Introduction

Understanding how functional neuronal circuits are established during development is a fundamental challenge in neuroscience. One prominent line of thinking is that network structure is initially only rudimentarily defined by genetic mechanisms, and that functional wiring is established through activity-dependent processes (Katz and Shatz 1996; Goodman and Shatz 1993; Hübener and Bonhoeffer 2014). Indeed, research in artificial intelligence and computational neuroscience have highlighted how weakly structured models can learn to perform complex tasks, even outperforming humans at Chess and Go (Silver et al. 2018, 2017). Such models are thought to abstract the biological phenomenon of experience-dependent rewiring of neuronal circuits, which is assumed to be critical for the maturation of the functional brain (van Gerven and Bohte 2018; Hasson, Nastase, and Goldstein 2020; Richards et al. 2019).

The significant role of sensory-driven neuronal activity in brain development was made apparent by the discovery of critical periods for ocular dominance formation in cats and primates (Hubel and Wiesel 1962, 1970; Hubel, Wiesel, and Stryker 1977). These seminal experiments, and many that followed, deprived animals of sensory inputs early in development and then showed major structural and functional abnormalities in the brain (Sengpiel, Stawinski, and Bonhoeffer 1999; Kind et al. 2002). In turn, theoretical and computational neuroscience work began to internalize the idea that stimulus-driven neuronal activity is sufficient, and at times necessary, for tuning the brain (Dayan and Abbott 2005). However, it is still unclear whether ‘normal’ sensory experience during development is indeed necessary for the emergence of complex behaviors.

More recently, the discovery of correlated spontaneous neuronal activity in early brain development has illuminated the potential for activity to tune neuronal circuits prior to the arrival of sensory input (Bajar et al. 2022; Ge et al. 2021). Indeed, wave-like activity patterns in the retina have been shown to contribute to the refinement of visual circuitry, and can even encode relevant statistical properties of natural stimuli (Ge et al. 2021; Warland, Huberman, and Chalupa 2006; Khona, Chandra, and Fiete 2022; Feller et al. 1996). However, perturbations of such developmental activity, including sensory deprivation studies, are usually restricted to small brain regions and target only a single sensory modality, while neighboring input streams remain intact. This generates competition between the perturbed modality and its unperturbed neighbors, making it impossible to distinguish whether a loss-of-function was caused by the lack of neuronal activity itself, or by competitive takeover from other modalities.

In summary, testing whether any, or all, of these activity-dependent components of development are truly necessary for the maturation of functional neuronal circuits requires the ability to reversibly block all neuronal activity throughout the period of brain formation, an intervention previously lethal at birth (Verhage et al. 2000; Sando et al. 2017). Here, we have overcome this challenge by pharmacologically blocking all neuronal activity patterns during the four days in which the central nervous system of the larval zebrafish is formed. We found that, after washout of the reversible agent, zebrafish could not only perform complex visuomotor behaviors, but they also exhibited fully functional and appropriately tuned neuronal cell types whose response properties were no different from those found in unperturbed animals.

## Results

The early maturation of functional circuits in larval zebrafish provides a unique opportunity to study the contributions of activity-dependent and -independent components of neuronal development for vertebrate behavior. For instance, when exposed to whole-field motion, six day old zebrafish can turn and swim to match the direction and speed of the visual stimulus, thereby allowing the animal to stay stationary in a moving stream (Roeser and Baier 2003; Neuhauss et al. 1999; Maaswinkel and Li 2003). This complex sensorimotor behavior, known as the optomotor response (OMR), is governed by multiple neuronal populations with distinct response properties and time constants that are activated across various brain regions (Fetcho and O’Malley 1997; Ahrens et al. 2012). Notably, their interplay encodes complex computations, ranging from motion discrimination (Naumann et al. 2016; Orger et al. 2008), to evidence integration (Seth, Prescott, and Bryson 2011; Harpaz, Aspiras, et al. 2021; Bahl and Engert 2020; Dragomir, Štih, and Portugues 2020), and swim-decision making (Harpaz, Nguyen, et al. 2021; Barker and Baier 2015). Thus, the OMR presents a rich paradigm for studying the contributions of sensory experience, neuronal activity, and activity-independent mechanisms in the maturation of a complex neuronal circuit.

In order to quantify the larval zebrafish’s OMR, we utilized a custom-designed behavioral rig (Bahl and Engert 2020) (Figure 1a), in which animals can be exposed to gratings that move orthogonally to their body axis (in either leftward or rightward direction) under closed-loop conditions (Naumann et al. 2016; Orger et al. 2008). When shown no stimulus, fish mainly swam forward (zero degree turns), but also made small-angle turns to the left or right, as seen in the “shoulders” around the dashed line at zero degrees (Figure 1b). However, when presented orthogonally moving gratings, animals displayed a strong tendency to enhance the probability of turning in the stimulus direction, which can be observed as a “bump” to the left of zero (dashed vertical line) for leftward moving stimuli, and to the right of zero for rightward moving stimuli (Figure 1c). This asymmetry in turn statistics disappeared for forward-moving stimuli. We can summarize these patterns of response to directed whole-field motion by plotting the average cumulative turn angle for all fish under the different stimulus conditions (Figure 1d), which highlights the persistent turn preferences over the course of an experiment. In addition to turn probabilities, we can also quantify the proportion of “correct” turns over all swim bouts, which provides a sensitive summary metric for behavioral performance that is used throughout the manuscript (Figure 1e). Lastly, we examined the overall swim activity of zebrafish larvae under all stimulus conditions, and found that their bout rate increased from 0.5 Hz in the absence of stimulation (Figure 1b inset) to 1 Hz when whole-field motion of any direction was presented (Figure 1c inset).

**Figure 1.**
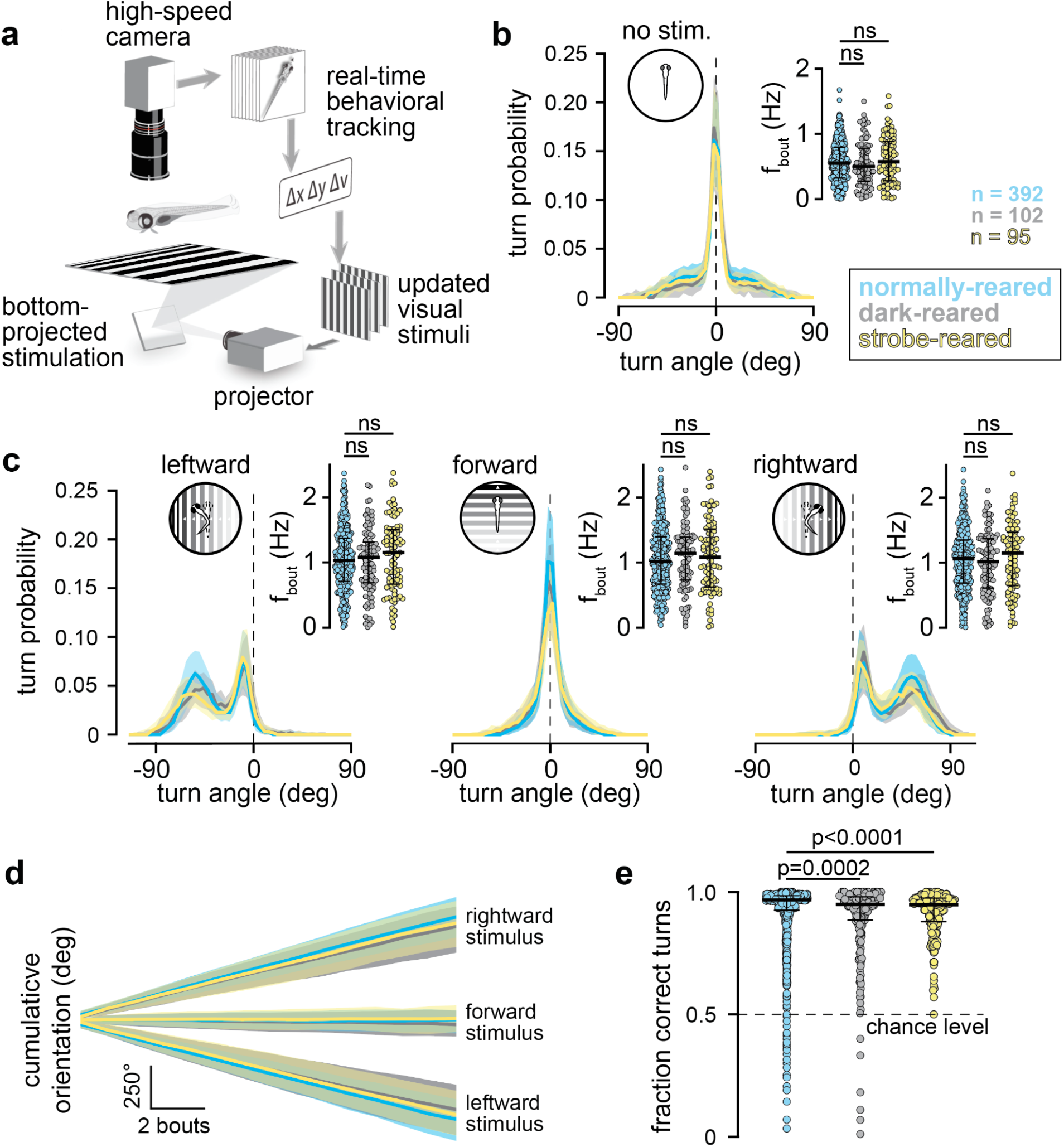
Visual experience is not necessary for the maturation of OMR. **(a)** Experimental setup; freely swimming fish are monitored with a camera while moving gratings are presented. Heading (Δν) and position (Δx, Δy) are extracted to lock visual motion direction to the body axis. Modified with permission from Naumann et. al. (2016). **(b)** Turn statistics of fish in absence of visual stimulus. Lines indicate median probability of swim bout angles for fish reared under a normal 14/10 hour light/dark cycle (blue), in total darkness (black), and for 14 hours in a 1 Hz strobe and 10 hours in darkness for the entirety of development (yellow). Shading indicates quartiles of the probability of performing a given swim bout angle. Inset: average bout frequency of individual fish under the same experimental conditions. **(c)** Turn statistics and bout frequencies of fish when shown leftward-moving (left), forward-moving (center), and rightward-moving gratings (right). Notation as described in (b). **(d)** Cumulative orientation change over time for control, dark-reared and strobe-reared fish. Lines and shading indicate median responses and quartiles for rightward-moving (top), forward-moving (middle), and leftward moving stimuli (bottom). **(e)** Proportion of correct turns made by control fish, dark-reared fish, and strobe-reared fish. Circles correspond to performance of individual animals over all trials (horizontal bar; median of correct turns across all fish).

In order to quantify the contribution of early sensory experience for OMR, we raised zebrafish in complete darkness, thereby eliminating visual experience altogether (Avitan et al. 2017). Generally, larval zebrafish start swimming immediately after they hatch at 2-3 days post fertilization (dpf), allowing them several days of visual and motor experience prior to our quantification of OMR at 6 dpf. Thus, our dark-reared animals swam and behaved for several days in the absence of any visual input. When we applied our behavioral quantification protocol to dark-reared larvae, we found no detectable difference in any of our metrics (“dark,” black vs “control,” blue), except for a small decrease in the fraction of correct turns in dark-reared animals (Figure 1b-e). We additionally considered whether irregular visual experience during development might have a deleterious effect on behavior, for which we raised fish under a constant 1 Hz strobe light during daytime (Rodger et al. 2006; Brickley et al. 1998). Yet, similar to the dark-reared individuals, strobe-reared fish showed no change in any of our behavioral metrics (yellow, Figure 1b-e), with the exception of a slightly smaller fraction of correct turns in the perturbed animals. These results suggest that natural visual experience was not necessary for the emergence of the OMR, prompting the study of earlier mechanisms for the maturation of this complex behavior.

Even in the absence of any sensory stimuli, spontaneous activity patterns, such as retinal waves, are thought to provide informative signals for the self-organization of neuronal circuits during development (Bajar et al. 2022; Martini et al. 2021; Guillamón-Vivancos et al. 2022). In the zebrafish visual system, such spontaneous activity in retinal ganglion cells (RGCs) begins at 2.5-3.5 dpf (Zhang et al. 2010, 2016), right before RGCs form functional projections to the optic tectum (by 4 dpf) (Burrill and Easter 1994; Stuermer 1988), illustrating a well-characterized synaptic pathway for visually-guided decisions that could be subject to activity-dependent maturation processes. To test to what extent spontaneous neuronal activity, across any and all circuit elements, is necessary for the maturation of functional neuronal circuits in larval zebrafish, we developed a protocol to raise zebrafish under complete anesthesia. Amongst a series of potential candidates that are used as anesthetics in fish and amphibia, we settled on tricaine, a sodium-channel blocker known to reversibly silence neuronal action potential firing (Popovic et al. 2012; Stanley et al. 2020; Ramlochansingh et al. 2014; Leyden et al. 2022).

We first measured the acute effect of tricaine on spontaneous and visually-evoked swimming behavior, and observed that all animals stopped moving within 10 seconds of tricaine immersion, at which point fish generally lost postural control and settled on the bottom of the dish (Supplemental Movie 1). Further, we found that fish resumed their swimming behavior within minutes of anesthesia washout, although with distinctly reduced vigor (Supplemental Movie 2).

In order to confirm tricaine’s function as a global and reversible blocker of neuronal activity, we performed functional calcium imaging experiments to quantify brain-wide responsiveness under anesthesia (Figure 2a). We began by imaging normally-reared zebrafish (1) before anesthesia, (2) during a one-hour tricaine treatment, and (3) after washout of the drug (Figure 2a,d). In this way, we could observe the degree to which neuronal activity is silenced under tricaine application and quantify the timescale and magnitude of recovery after anesthetic washout.

**Figure 2.**
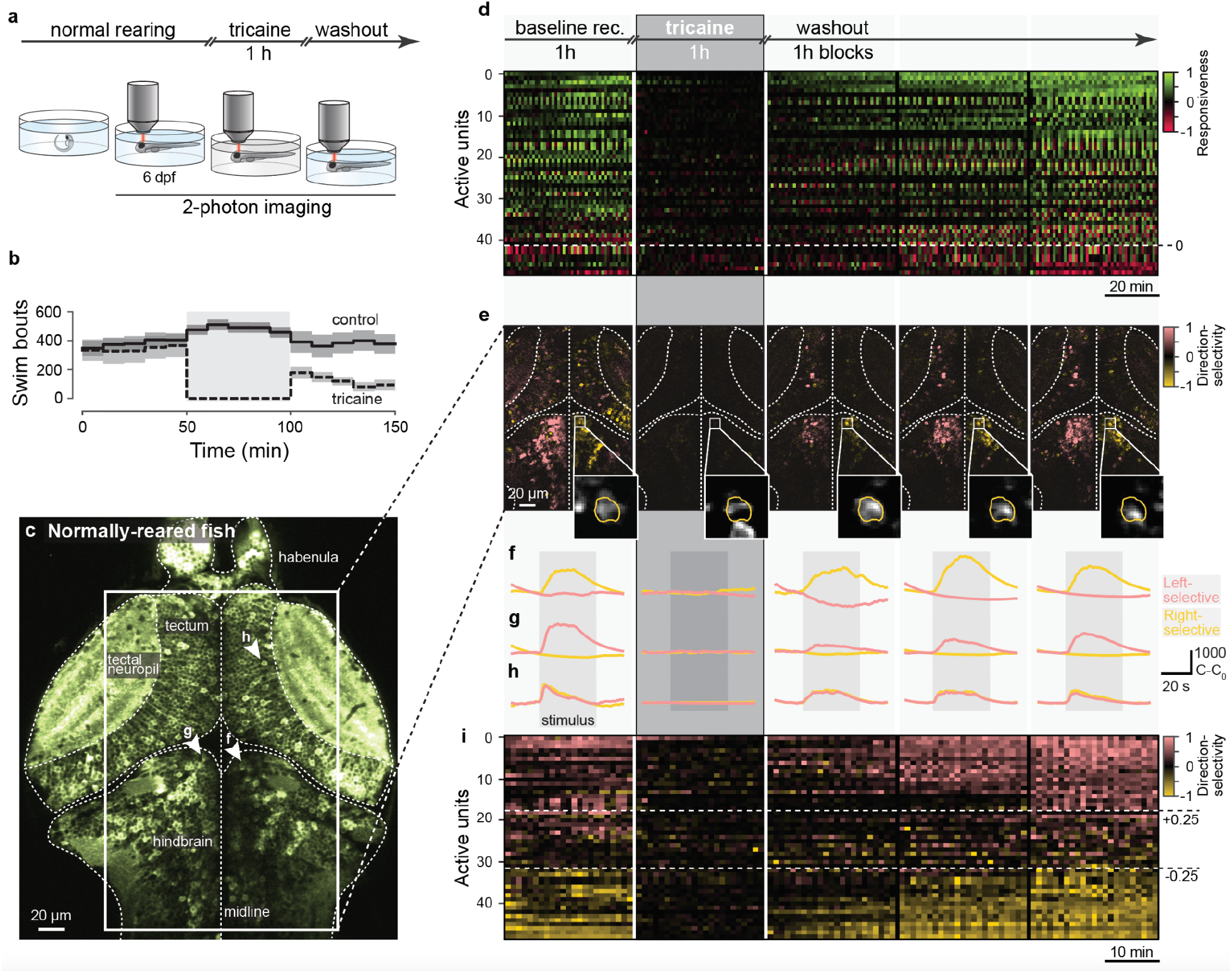
Tricaine anesthesia silences neuronal activity reversibly. **(a)** Experimental paradigm: fish are raised under standard conditions; at 6 dpf, we embedded and imaged calcium activity for 1 h, imaged under tricaine for 1 h, and finally imaged as neural activity returned during tricaine washout. **(b)** Effect of tricaine anesthesia on the number of bouts during OMR. Solid line, awake fish raised under control conditions; dashed line, fish raised under control conditions and anesthetized for 1 h (shaded region). Lines, mean bout rate; gray bands, standard deviation. **(c)** Averaged calcium signal prior to tricaine administration including regions involved in the visuo-motor processing hierarchy (indicated). The overlaid box indicates the region analyzed in d-i; arrowheads indicate cells shown in f-h. **(d) Top**, schematic showing time-course of calcium imaging, each block contains 1 h of recording. Remainder of the figure is aligned to imaging blocks. **Bottom**, responsiveness-index of units in tectum, pretectum, and hindbrain (see Methods; 3 fish). Units above the dashed line on average increase activity in response to stimuli, units below on average decrease activity during stimulus presentation. Note strong responsiveness levels during baseline return by the end of the washout period. **(e)** Leftward (pink) and rightward (yellow) direction-selectivity index (see Methods) of region highlighted in c. Note that strong left and right direction-selectivity during baseline disappears during anesthesia and slowly recovers during washout. Insets show a rightward-selective example neuron (yellow outline) that was imaged across the entire experiment (corresponds to traces in f); each image normalized separately. **(f-h)** Averaged 1-minute trials showing stimulus-evoked activity of a right-selective (f), left-selective (g), and motion-selective (h) cell during baseline, tricaine, and washout periods. Note that visual responsiveness disappears during tricaine and that the same direction-selective tuning re-emerges during washout. **(i)** Direction selectivity-index of units in the tectum, pretectum, and hindbrain (see Methods; 3 fish). Units above the top dashed line are distinctly left-selective, while units below the bottom dashed line are distinctly right-selective. Note that the direction-selectivity during baseline returns by the end of the washout. Our algorithm to compute direction-selectivity has a unit-wise normalization that is sensitive to very low activity levels (see Methods), thus resulting in noisier signal during tricaine. Units in d, e, i consist of 1-5 neurons with same response properties during visual stimulation, thus representing small computational circuit blocks.

We first measured the effect of tricaine on visually-evoked activity across several brain areas known to be involved in processing visual information, including the tectum, pretectum and anterior hindbrain (Figure 2c). We found that anesthetic application dramatically reduced neuronal activity from baseline within minutes (Figure 2d) and that activity recovered over a period of 2-3 hours after washout in all tested animals (Supplemental Figure 1), results that are in general agreement with the observation that extended recovery times were necessary to restore fish to full behavioral vigor after similar treatment (Figure 2b).

We next extended our analysis to the response properties of individual units, where we focused on neuronal response types that are critical for the OMR. Globally, we observed distinct left-selective and right-selective neuronal populations (Figure 2e), which became unresponsive under tricaine application and gradually returned to baseline responsiveness levels in the hours after anesthesia washout. To quantify the extent of anesthesia and recovery, we identified individual rightward-selective, leftward-selective and motion-selective cells that we were able to record from for the entire duration of the experiment (Figure 2f-h) and observed how their response properties changed during anesthesia and washout. We found that, similar to global activity patterns, specific response properties disappeared with the onset of anesthesia and returned to baseline gradually after removal of the block (Figures 2f-h). Figure 2i highlights this phenomenon by showing the direction-selectivity of many units over individual trials throughout the five hours of the experiment. Here, it again becomes apparent that anesthetic application dramatically reduced neuronal activity from baseline already by the first visual-stimulation trial, and that activity gradually recovered over the following hours of washout.

Having validated the efficacy of tricaine as a reliable and reversible anesthetic for the larval zebrafish brain, we utilized the drug to block all neuronal activity from 36 hours post fertilization until behavioral testing at 6 dpf. This four-day period of neuronal inactivation includes all critical phases of circuit maturation and starts well before the emergence of spontaneous neuronal activity in larval zebrafish at 2-3 dpf (Zhang et al. 2010, 2016; Burrill and Easter 1994; Stuermer 1988). While many fish raised in anesthesia displayed normal development, we found that some had anatomical deficits, including uninflated swim bladders and hunched backs (Figure 3a). Informed by the multi-hour recovery of neuronal activity we observed after the relatively brief one-hour tricaine exposure described above (Figure 2, Supplemental Figure 1), we performed behavioral testing immediately after anesthetic washout, as well as two, six and 24 hours after release from anesthesia. We found that even immediately after tricaine removal, all anesthesia-reared fish could swim, as indicated by a non-zero bout rate under all stimulus conditions (Figure 3b) and they could see, as evidenced by an increase in bout rate when presented with forward-moving gratings (Figure 3b). Further, tricaine-reared fish were capable of computing the direction of whole-field motion immediately after washout, as their turning probability was biased towards the ‘correct’ stimulus direction (Figure 3c, 0-1 hour, median of 55% accuracy of turns). Over the subsequent periods, performance consistently increased, reaching 75% median accuracy by 2 hours of washout, 89% median accuracy by 6 hours, and 93% by 24 hours, comparable to the 96% observed in normally-reared individuals (Figure 3c).

**Figure 3.**
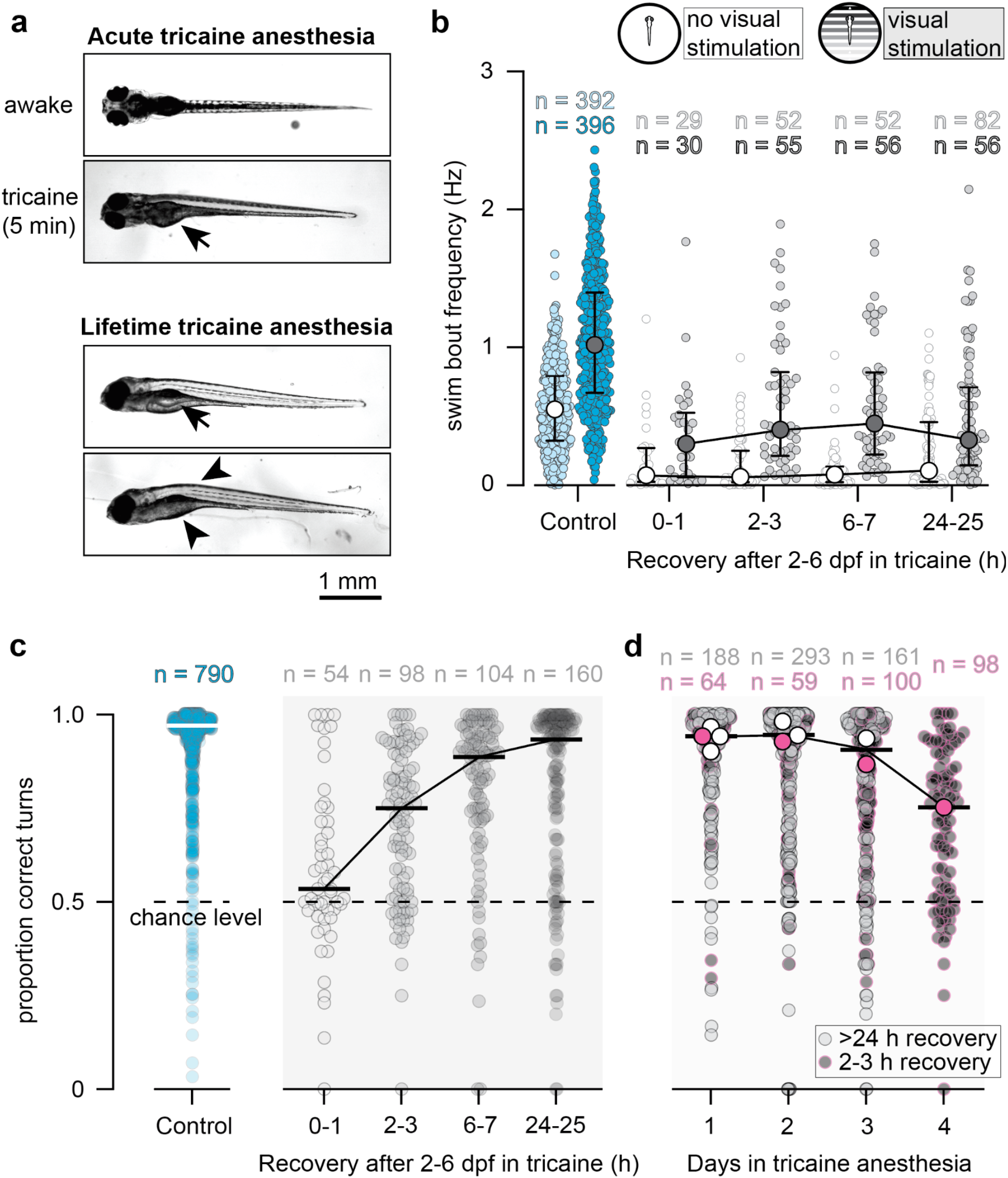
Fish raised under complete anesthesia can see, swim, and integrate visual stimuli. **(a) Top**, within 1 minute of acute tricaine application, zebrafish lose their ability to swim (Supplementary Movie 1) and to maintain postural control, rolling onto their sides; arrow, inflated swim bladder. **Bottom**, adverse morphological effects of raising fish in anesthesia animals cannot inflate swim bladders (arrow), as they do not swim to the surface to gulp air. Occasionally, we observed hunched backs and abnormal abdominal morphology (arrowheads). **(b)** Bout frequency of control fish (n = 396) and fish raised in anesthesia from 2-6 dpf; no visual stimulation (light blue, white) and during OMR (dark blue, gray) after varying amounts of recovery time post anesthetic washout. Note that under all conditions, except during tricaine, fish increase their bout frequency when shown forward moving gratings compared to no stimuli. **(c)** Proportion of correct turns during OMR made by control fish (blue, left) or fish raised under anesthesia from 2-6 dpf. Groups were tested after a varying number of hours of recovery after washout (gray, right). Circles, individual fish’s performance over all trials; horizontal bars median of correct turns of all fish within a condition. **(d)** Proportion of correct turns during OMR made by fish raised under anesthesia for varying numbers of days during development. Circles, individual fish’s performance over all trials; horizontal bars; median performance over all n-day treatments; open circles, median values for each possible condition (e.g., ‘1 day’ includes fish placed in tricaine for 24 hours starting on either days 2, 3, 4, or 5 of development, ‘2 day’ includes fish placed in tricaine for any 48 hour-period starting on either days 2, 3 or 4 of development, and hence contain 4 and 3 open circles, respectively). All fish that were taken out of anesthesia on the day of testing received 2 hours of washout (pink circles), therefore the ‘4 days’ group in (d) corresponds to ‘2-3 hours’ group in (c); all other fish recovered for >24 h before testing.

We next tested the effects of shorter tricaine-treatment periods on behavioral performance. To that end, we anesthetized fish for several possible one-, two-, or three-day periods during the first six days of their development (Figure 3d). Most treatment cohorts were allowed to recover for at least one day or more from the anesthetic before behavioral testing (Figure 3d, gray), with the exception of four groups, for which fish were left in the anesthetic up until the time of behavioral testing and recovery was limited to two hours (Figure 3d, pink). While fish undergoing four-day tricaine treatment required six hours of recovery to reach full behavioral performance, we found that shorter periods of anesthesia had only marginal effects on OMR performance: no single-day, two-day or three-day treatment led to a drop below 90% accuracy, even when measured after a recovery period of only 2 hours (Figure 3d).

One explanation for the significantly extended recovery time after four days of anesthesia could be a compromised general physiology caused by the extended tricaine exposure itself. These fish were not only deprived of brain activity throughout development, they also never swam or utilized their muscles until washout, which in itself has been shown to cause decreased cell proliferation in the larval zebrafish forebrain (Hall and Tropepe 2018). Indeed, we noticed distinct anatomical differences in the brains of tricaine-reared animals (Figure 4a,b) when compared with normally-reared animals (Figure 4c), which included an overall smaller brain-cross section, larger ventricles, less densely packed cell bodies and neurons with bright fluorescence signals that could indicate ongoing apoptosis. In addition, chronic tricaine exposure had adverse effects on zebrafish body development, including swim bladder defects, reduced body size, and generally abnormal morphology (Figure 3a).

**Figure 4.**
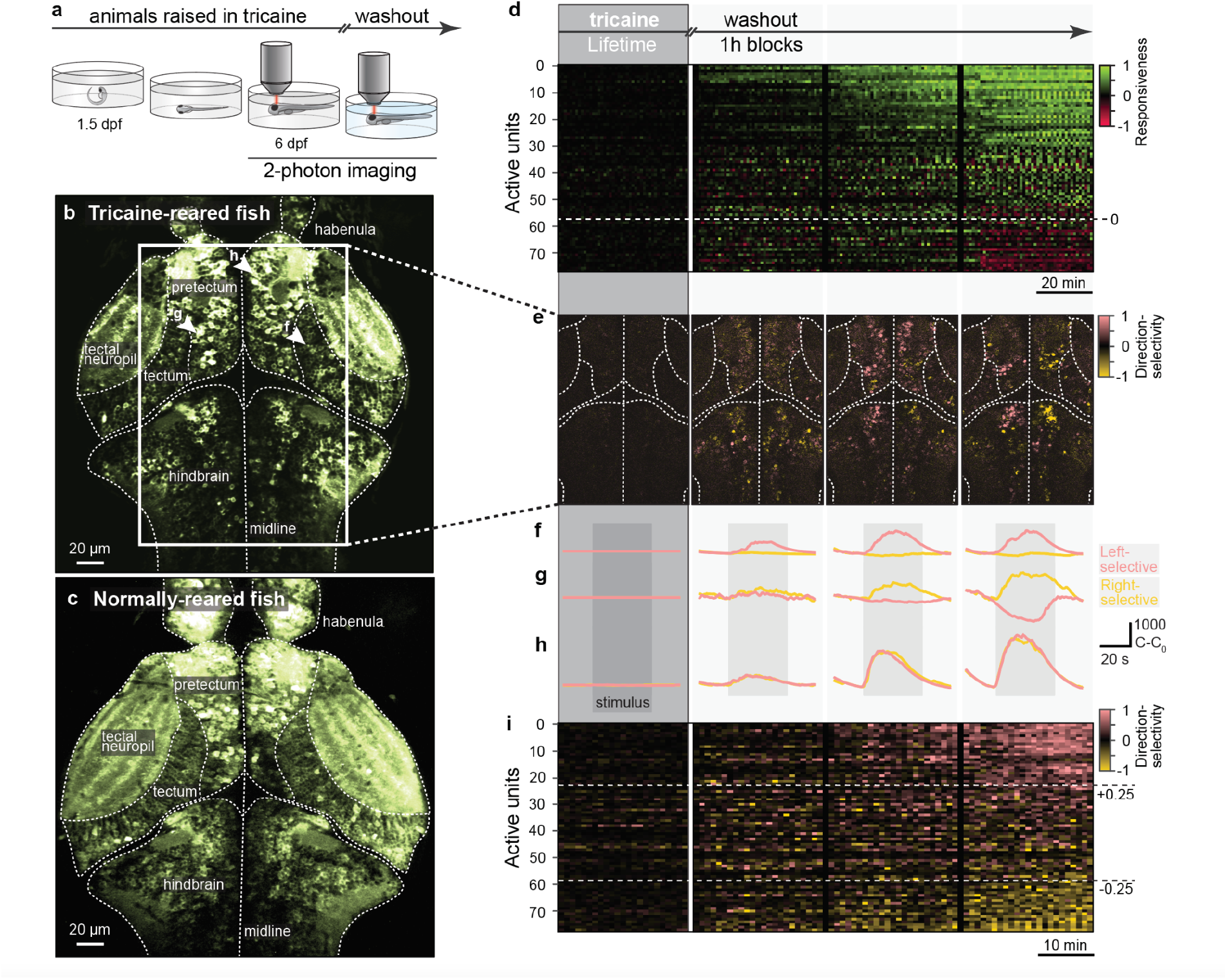
Tricaine-reared fish display fully tuned neuronal responses after anesthetic washout. **(a)** Experimental paradigm: fish are raised under anesthesia from 1.5 dpf onwards; at 6 dpf, we embedded and imaged calcium activity under tricaine for 1 h, then imaged as neural activity returned during tricaine washout. **(b)** Averaged calcium signal in a tricaine-reared fish following anesthetic washout. The overlaid box shows the region analyzed in (e), arrowheads indicate cells shown in f-h. Noteworthy morphological aberrations in tricaine-reared fish included overall smaller brain cross-section, extended ventricles, smaller neuropil regions, sparser cell populations, and exceptionally bright neurons that could indicate ongoing apoptosis. **(c)** Averaged calcium signal in an awake, normally-reared fish; similar imaging plane as in b for anatomical comparison. **(d) Top**, schematic showing time-course of calcium imaging, each block contains 1 h of recording. Remainder of the figure is aligned to imaging blocks. **Bottom**, responsiveness-index of units in tectum, pretectum, and hindbrain (see Methods; 3 fish). Units above the dashed line on average increase activity in response to stimuli, units below on average decrease activity during stimulus presentation. Note strong responsiveness levels emerge by the end of the washout period, comparable to normally-reared animals (Figure 2). **(e)** Leftward (pink) and rightward (yellow) direction-selectivity index (see Methods) of region highlighted in c. Note that strong left and right direction-selectivity are absent during anesthesia and slowly appear during washout. **(f-h)** Averaged 1-minute trials showing stimulus-evoked activity of a right-selective (f), left-selective (g), and motion-selective (h) cell during tricaine and washout periods. Note that visual responsiveness is absent during tricaine and that direction-selective tuning emerges during washout. **(i)** Direction selectivity-index of units in the tectum, pretectum, and hindbrain (see Methods; 3 fish). Units above the top dashed line are distinctly left-selective, while units below the bottom dashed line are distinctly right-selective. Note that direction-selectivity appears by the end of the washout, comparable to normally-reared animals (Figure 2). Our algorithm to compute direction-selectivity has a unit-wise normalization that is sensitive to very low activity levels (see Methods), thus resulting in noisier signal during tricaine. Units in d, e, i consist of 1-5 neurons with same response properties during visual stimulation, thus representing small computational circuit blocks.

An alternative explanation for the extended recovery time after our four-day anesthetic protocol is that functional neuronal circuit development requires several hours of compensation by activity-dependent tuning. That is, fish reared under anesthesia might not have an innate ability to perform OMR, but rather require a block-free period in order to rewire the circuit after tricaine washout. To distinguish between these two alternatives, we next set out to perform brain-wide imaging experiments in animals that emerge for the first time in their lives from tricaine anesthesia (Figure 4a).

As before (see Figure 2), we measured visually-evoked activity across brain areas known to be involved in processing visual information (Figure 4b), (1) during continued tricaine treatment and (2) after washout of the drug (Figure 4a,d). Reassuringly, during the first hour of recording (while animals remained anesthetized) no responsiveness to visual stimulation could be seen (Figure 4d, leftmost panel) and overall fluorescence was essentially absent (Figure 4e, leftmost panel), thus confirming a complete block of activity by tricaine anesthesia (Supplemental Movie 3). Next, we identified individual neuronal response types, such as motion- and direction-selective neurons, whose role is well-characterized in the OMR circuit (Naumann et al. 2016) (Figure 2). Such cell types might first become active in an unspecific fashion and only acquire their tuning properties gradually, indicating a need for activity-dependent mechanisms throughout development. Alternatively, cell types might appear immediately after washout in a fully-tuned fashion, which would suggest that the network developed into a functionally-tuned state in the absence of any neuronal activity and just needs to be relieved from anesthesia.

Remarkably, we found that the responsiveness and direction-selectivity tuning of the visual circuitry emerged from anesthesia at equivalent rates in animals that were anesthetized for only an hour (Figure 2d,e,i) or for the entire duration of development (Figure 4d,e,i). Moreover, we found that the different direction-selectivity preferences of individual neurons, from which we were able to record for the entire duration of the experiment, also emerged at the same rate and in the same gradual nature when compared to fish that were anesthetized for only one hour (Figure 2e-g vs Figure 4e-g). Thus, we conclude that the emergence of functionally-tuned circuits in tricaine-reared fish does not depend on neuronal activity-based rewiring during washout. Rather, we suggest that functional response types are established by activity-independent mechanisms and that the washout period serves solely to remove the damping effects of anesthesia.

In summary, our results suggest that the emergence of the neuronal circuitry for seeing, swimming, and sensory-motor integration is remarkably independent of developmental neuronal activity. Instead, we propose that many of the computational building blocks of the underlying functional circuitry, including responsiveness to visual stimuli, direction-selectivity, motor command neurons and central pattern generators in the spinal cord develop in an activity-independent manner under the exclusive control of genetic and transcriptional algorithms that are read out from the genome of the animal (Kovács, Barabási, and Barabási 2020; Barabási and Barabási 2020).

## Discussion

Here, we have developed a reversible lifetime block of sensory stimuli and neuronal activity in a vertebrate that, when removed, reveals a functional and appropriately-tuned brain. In this way, we were able to separate “nature” and “nurture” for the first six days of an animal’s life, thereby providing a toolkit for studying the extent of “innate” neuronal wiring in a vertebrate model system. Using this approach, we demonstrate the remarkable extent to which activity-independent developmental mechanisms precisely pattern neuronal circuits.

We believe that our results contextualize, rather than overturn, a long literature of developmental perturbations, including eye sutures, sensory deprivations, and genetic silencing of activity that have suggested a crucial contribution of neuronal activity for proper brain development. We note that these experiments generally relied on regional (and not global) plasticity perturbations and, critically, they introduced a competitive imbalance between different modalities or mixed input channels. Amongst the most prominent examples are the monocular occlusion experiments by Hubel and Wiesel, where a clear competitive advantage was introduced to the eye that was allowed to remain open (Hubel and Wiesel 1970; Wiesel and Hubel 1963). Interestingly, in less popular work, Hubel and Wiesel sutured both eyes shut prior to eye-opening, and nevertheless found the visual cortex in a remarkably unperturbed state (Wiesel and Hubel 1965), a result recently reproduced in monkeys (Arcaro, Schade, and Livingstone 2018). Even the remaining aberrations of visual cortex neurons can be explained by the fact that perturbed visual inputs are outcompeted by other, non-visual modalities, such as motor-related or auditory signals, which have been demonstrated to contribute to primary visual cortex processing (Keller, Bonhoeffer, and Hübener 2012; Garner and Keller 2022). Another impressive example of competitive takeover of underused cortical regions is the finding that visual pathways can successfully innervate auditory cortex in ferrets when the afferent thalamic relay nuclei are surgically removed (von Melchner, Pallas, and Sur 2000; Sharma, Angelucci, and Sur 2000). This phenomenon is not restricted to the sensory cortex: in the mouse olfactory bulb, it has been shown that blocking an individual olfactory channel leads to significant rearrangement of synaptic connectivity, while blockage of all receptors leaves the circuitry largely unchanged (Yu et al. 2004).

Another experimental perturbation that was found to interfere with circuit refinement is the local application of synaptic transmission blockers to interrupt afferent neuronal inputs. For example, it was found that N-methyl-D-aspartate (NMDA) receptor antagonists, when locally applied to the optic tectum of frogs (Cline and Constantine-Paton 1989) and rats (Simon et al. 1992) prevented pruning of incoming axonal arborizations from RGCs. Similarly, local curare application to the neuromuscular junction in mice is known to prevent the segregation of motor neuron axons onto individual myofibers (Sanes and Lichtman 1999; Tapia et al. 2012). Classically, these results support the notion that activity-dependent processes are necessary for the refinement of circuit structure. Yet, these perturbations categorically block trans-synaptic, homeostatic signaling cascades, which are also known to play a critical role in the signaling between pre- and postsynaptic elements in early development, including target recognition and functional stabilization. Thus, these results can not distinguish whether spike timing-dependent plasticity or synaptic signaling of any sort is responsible for the observed refinement of circuit structure.

Therefore, our hypothesis is that the default innervation of brain regions is governed exclusively by genetic programs and does not depend on competitive interactions. This predicts that if competition is removed by a categorical block of all neuronal spiking activity, as was done in our experiments, the ‘natural,’ hard-wired brain structure should emerge, in spite of this rather drastic perturbation. Indeed, in very early tadpole experiments it was revealed that normal behavioral patterns can be observed in cloretone-reared animals, a treatment that was assumed to block all neuronal activity (Haverkamp and Oppenheim 1986; Harrison 1904). Later studies in developing mouse embryos have yielded similar results by showing that genetically blocking synaptic release in all neurons led to an anatomically unchanged macro-, micro- and nano-structure throughout the brain (Verhage et al. 2000). Moreover, in newborn kittens, the binocular features of the visual cortex were shown to emerge even if eye-specific, monocular visual stimuli were explicitly decorrelated from each other (Gödecke and Bonhoeffer 1996), suggesting that the emergence of functional structure in the cat visual system is governed largely by genetic programs, in a similar fashion to what we observed in the visual pathways of the larval zebrafish. As such, the application of global and reversible anesthetics allows to highlight and extract the prominent contribution of nature over nurture in early behaviors.

From an evolutionary perspective, a robust hardcoding of essential, basic behaviors is expected to be present in all animals. This can be seen in turtles heading out to sea after hatching (Carr, Ogren, and American Museum of Natural History 1960), newborn zebras galloping alongside their herd, and freshly-hatched iguanas escaping from snakes (Greene et al. 1978). As such, we propose that a large fraction of adaptive motor sequences and sensorimotor reflexes may be a result of hard-wired and innate circuitry. By contrast, environmental changes that cannot be anticipated on evolutionary timescales require explicit learning and adaptation to develop novel behavioral modules (Zador 2019). Nevertheless, given that complex behaviors can certainly emerge from activity-independent developmental processes, open questions remain as to (1) which behaviors and circuits do require neuronal activity for maturation and (2) why activity is utilized for refinement of neuronal circuits, when developmental processes could be sufficient. Further understanding and incorporating these neurodevelopmental processes may prove instructive to both computational neuroscience and machine learning communities in the future (Barabási, Beynon, and Katona 2021; Koulakov et al. 2021; Barabási and Czégel 2021).

## Acknowledgements

We thank Andrew Bolton for helping conceive the study and for assistance in pilot experiments. This work was supported by NIH NIGMS T32 GM008313 Grant to D. Barabasi, by Swiss National Science Foundation Postdoc Mobility Fellowships P2EZP3 188017 and P500PB_203130 to G. Schuhknecht, and by NIH Grant U19NS104653, NIH Grant 1R01NS124017, NSF Grant IIS-1912293, and Simons Foundation SCGB 542973 to F. Engert.

## Author Contributions

DLB and FE conceived the project. DLB developed the anesthetic protocol, performed behavioral experiments, and analyzed resultant data. GFPS performed two-photon imaging experiments. DLB and GFPS analyzed imaging data. All authors contributed to preparing figures and writing the manuscript.

## Data and Code Availability

The code used to generate figures is available at https://github.com/bdanubius/InnateFish. The repository includes a link to download processed data used for the figures, raw imaging data available at reasonable request from the corresponding author.

## Methods

### Zebrafish

For all behavioral experiments, we used 6 dpf wild-type (WIK) zebrafish. Fertilized eggs were collected in embryo water (Methylene Blue + E3 solution (5 mM NaCl, 0.17 mM KCl, 0.33 mM CaCl2, 0.33 mM MgSO4)) and transferred to filtered fish facility water after 24 h. Under standard rearing conditions, zebrafish were maintained on a 14 h light /10 h dark cycle at 28.5°C. Dark-reared embryos were placed in a light-proof box after collection, ensuring a 24 h dark cycle. After collection, strobe-reared embryos were placed in a box that contained a 1 Hz strobe light, which was active during the 14 h light period (matched to the standard rearing condition’s light on and off times) and off during the 10 h dark cycle.

All imaging experiments in this study were performed on the transgenic zebrafish line *elavl3:GCaMP6s* line (Bahl and Engert 2020). We raised groups of 20-30 larvae in standard Petri dishes (90 mm diameter) containing filtered fish water on a 14 h light, 10 h dark cycle at 28 °C. Imaging was at 6 dpf, at which age sex cannot be determined. Prior to imaging, we screened larvae for strong GCaMP6s fluorescence and the absence of any pigmentation. All experiments were approved by the Harvard University standing committee on the use of animals in research and training.

### Behavior experiments in freely swimming larval zebrafish

We performed free-swimming behavioral quantification with the experimental setup previously described in (Bahl and Engert 2020). Briefly, individual larvae were placed in a closed-loop virtual-reality environment and presented with a moving bar stimulus projected from below (60 Hz, AAXA P300 Pico Projector). In ‘no stimulus’ conditions, only a gray background was shown. We used real-time behavioral tracking at 100 Hz to determine the location and body orientation of animals and updated the visual stimuli accordingly.

Each behavior trial began with a 5s period in which gratings (spatial period of 1 cm and oriented along the animal’s anterior-posterior axis) were static and locked to fish orientation. These orientation-locked gratings began moving at a velocity of 1 cm/s for 10s, followed by another 5 s static period before the next trial was initiated. Each animal was presented with 30 sets of ‘no stimulus’ or forward-, backward, leftward-, or rightward-moving gratings. The order of stimuli presented within each set was randomized.

### Tricaine anesthesia

For all experiments, we used a concentration of 60 µg/ml tricaine in filtered fish water (MS-222 Sigma-Aldrich). We came to this concentration through extensive testing of a wide range of concentrations. We optimized for a solution that kept fish fully anesthetized for the maximum treatment duration of 4 days, while minimizing physiological effects and mortality. The tricaine bath was exchanged every 12 h because we found that tricaine started to lose its potency after around 18 h, presumably because it was metabolized by the animals kept in the dish (Supplemental Figure 2). We assessed whether larvae were indeed fully anesthetized through shaking the plate and tapping fish, as well as a long-term quantification of movement response, or lack thereof, under a light-on and light -off stimulus (Supplemental Figure 2). We discarded all fish in a petri dish if we found any animal that responded to visual or physical stimulus.

Zebrafish embryos were placed in petri dishes (9 cm diameter) filled with tricaine solution and kept in the dark until experimental testing, because tricaine forms toxic byproducts under extended exposure to light. Since the onset of spontaneous neuronal activity in larval zebrafish happens prior to hatching, we used a pronase solution (50 mg/ml in fish water for 5 min, Sigma-Aldrich) to dissolve the chorion prior to 24 hours post fertilization (hpf). This allowed us to anesthetize animals at 36 hpf and to have animals develop fully in the absence of their ability to hatch themselves.The same pronase-treatment was also performed for all ‘triaine-control’ fish used in the study.

### Two-photon calcium imaging

For *in vivo* imaging experiments, we prescreened larvae as described above. Larvae were then fully embedded in lukewarm agarose, which was allowed to solidify for one hour. We found that incorporating this resting period almost completely reduced drift of the imaging plane during recording sessions. Critically, this allowed us to record from the same identified units for extended periods of time.

Fish that were raised under tricaine anesthesia were embedded in an agarose solution containing 60 µg/ml tricaine to ensure that the embedding process would not relieve the anesthetic block. We used a custom-built two-photon microscope containing a femtosecond-pulsed MaiTai Ti:Sapphire laser (Spectra Physics) tuned to 950 nm for GCaMP6s imaging, a set of x/y-galvanometers (Cambridge Technology), a 20x obJective (XLUMPLFLN, Olympus), and a photomultiplier that was amplified by a SR570 current preamplifier (Stanford Research). The setup was operated by a custom-written Python 3.7-based software package. We tuned the laser power at the specimen to ∼13 mW and acquired images at frame rates of 1 Hz.

We imaged each plane at a spatial resolution of ∼0.7 µm/pixel (700 × 700 pixels) for one hour, during which animals were presented with a bottom-projected visual stimulation paradigm. Each visual stimulation trial lasted for 60 seconds and consisted of a 10-second period during which a stationary grating was presented, a 30-second period during which the grating then moved at constant velocity either to the left or the right of the animal (in a randomized fashion), and finally a 20-second period during which the grating was stationary again.

We designed a custom-built gravity pump system to exchange the liquids surrounding the embedded animal in the 2-photon microscope. Critically, this enabled us to replace the fish water surrounding the animal with tricaine solution (and vice-versa) without opening the setup or moving the objective, thus allowing us to image the same units for several hours under different anesthetic conditions (Figures 2, 4).

### Preprocessing of two-photon imaging data

We processed our imaging data with the open-source toolbox CaImAn (Giovannucci et al. 2019) and the Computational Morphometry Toolkit (Rohlfing and Maurer 2003), as described before (Bahl and Engert 2020). Briefly, we implemented a three-step processing pipeline, in which we first performed piecewise rigid motion correction using CaImAn (NoRMCorre; using standard parameters) (Rohlfing and Maurer 2003; Pnevmatikakis and Giovannucci 2017).

For the single-unit analyses shown in Fig. 2 f-h and Fig. 4 f-h, we applied CaImAn’s segmentation algorithm (CNMF; using standard parameters adjusted for frame rate and cell size). We used a relative measure for calcium dynamics by subtracting the baseline calcium level (C_0_; taken in the 10 s period prior to visual motion onset) from the calcium activity during the entire trial (C). We used this C - C_0_ metric instead of computing /1C/C_0_, because C_0_ was often zero. Importantly, for all single units analyzed in this paper, we confirmed visually that we indeed recorded from the same individual neurons during the entire duration of the experiment (i.e., a total of 5 hours) and excluded units that moved in or out of the imaging plane during the experiments.

For the multi-unit analyses shown in Fig. 2 d & i and Fig. 4 d & i, we hand-segmented small regions containing multiple neurons in motion-corrected image stacks. These were selected such that they contained clusters of 10 - 20 neurons that displayed the same functional responses during visual stimulation. Thus, these units correspond to small, similarly-tuned circuit motives across the tectum, pre-tectum, and hindbrain.

The responsiveness index was calculated by computing the average C-C_0_ value for each trial and by normalizing these values across all trials for each unit individually. Importantly, by normalizing each unit to its own maximum C-C_0_ value in this manner (as opposed to a ‘global’ normalization), we sought to highlight subtle responsiveness levels, particularly during tricaine periods, as an additional confirmation that brain activity was indeed silenced by anesthesia. Units that showed a net increase in responsiveness to the visual stimulation were normalized to 1, while units with a net decrease in responsiveness during visual stimulation were normalized to -1. This allowed us to dissect how tricaine anesthesia influenced excitatory and inhibitory circuit properties, respectively.

To compute the direction selectivity index, we matched subsequent pairs of trials in which gratings were first moving leftwards, followed by rightwards. We then calculated the difference in fluorescence responses to these two stimuli, which, after normalization, gave us a metric that ranged from -1 (i.e., rightward-selective) to +1 (i.e., leftward-selective). Given that we normalize each cell’s activity, we encounter a higher level of noise in the tricaine period, as low neural activity becomes indistinguishable from a lack of direction selectivity.

