## Supplemental Figures for "Nature over Nurture: Functional neuronal circuits emerge in the absence of developmental activity"

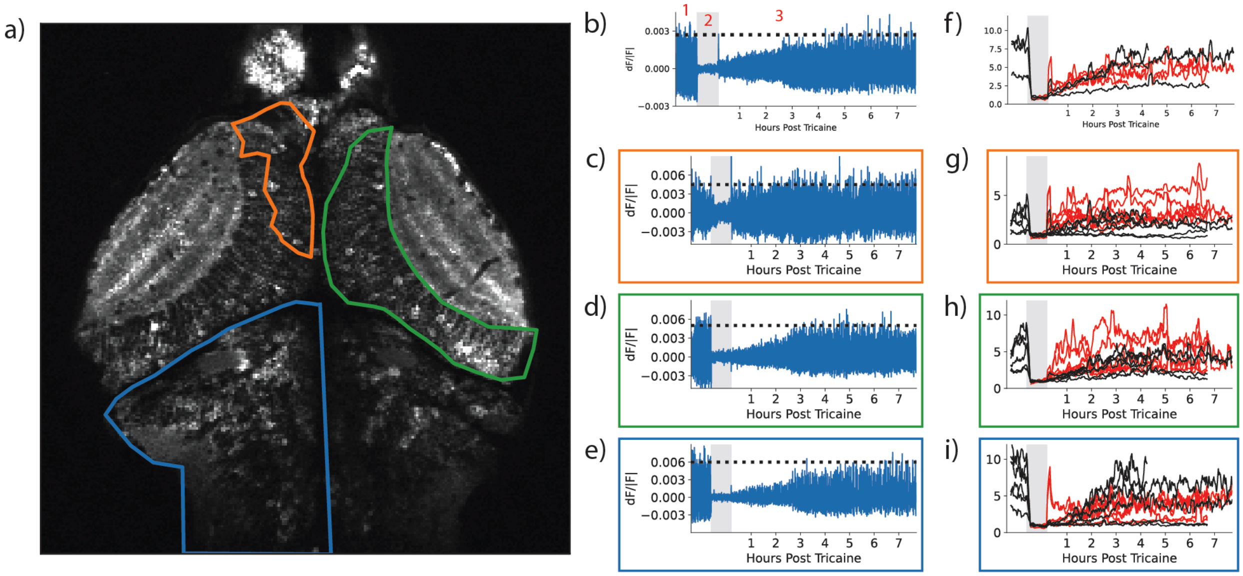

**Supplemental Figure 1**

**(a)** Average image of neural activity recording prior to tricaine administration of fish in Figure 2. Highlighted is the left hindbrain (blue), right tectal cells (green) and left pretectum (orange).

**(b)** Change in fluorescence (dF/|F|) of the whole imaging plane (1) during baseline, (2) during tricaine administration (2, gray region), and (3) after tricaine washout for the fish in (a). Dashed line indicates average activity before tricaine administration. Application of anesthesia leads to a categorical decrease in evoked activity. Activity increases immediately after washout, but takes 4-6 hours for a full recovery to baseline levels.

**(c-e)** Evoked activity (dF/|F|) of (c) pretectum, (d) tectum, and (e) hindbrain before, during and after tricaine application. All show a significant decrease in activity under tricaine application, but have different rates of recovery: pretectum (c) recovers to baseline immediately after washout in this fish, tectum (d) takes up to 3 hours for a full recovery, and hindbrain (e) can take up to 7 hours to recover to baseline levels.

**(f-h)** Evoked activity for multiple fish of the entire imaging plane (f), pretectum (g), tectum (h), and hindbrain (i). Black traces correspond to fish raised normally and placed under tricaine for 1 hour, while red traces indicate fish that were raised in anesthesia and had tricaine washed out for the first time during imaging. The timescales of recovery seem overall identical independent of the length of anesthetic application, supporting the results of Figures 3 and 4. Over all fish activity recovers rapidly in the first 2-3 hours of washout, then seems to gradually to a maximum level by 6 hours, supporting the 0, 2 and 6 hour timepoints of behavioral testing in Figure 3. In these subpanels we are plotting the upper envelope of the dF/|F| shown in (b-e), normalized to the tricaine period.

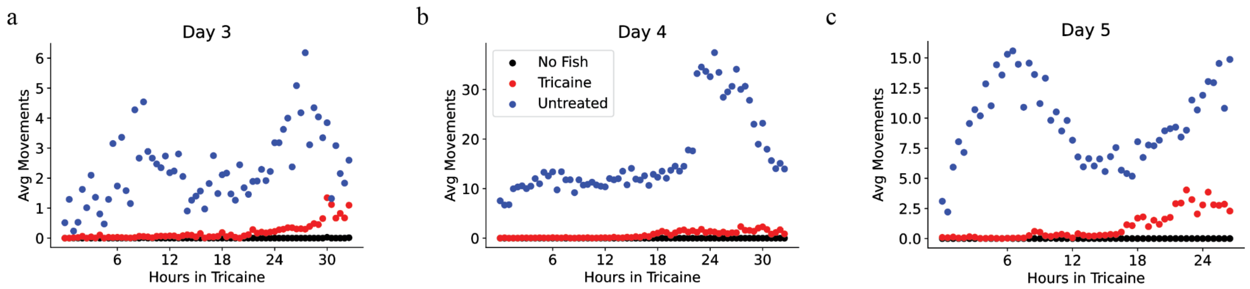


**Supplemental Figure 2**

(a-c) Quantification of the half-life of anesthesia treatment at various days of maturation. Fish are placed in 96 well plates in the dark at 3 days (a), 4 days (b) and 5 days (c) of age, prior to which they are kept in anesthesia. During the experiment, fish are shown a 30s light on and then a 30s light off stimulus every 10 minutes, and we record resultant motion. In all three cases we see clear motion in untreated fish throughout the experiments, while we see little to no motion in fish raised in tricaine prior to 18 hours of treatment. For this reason, we exchange the tricaine bath of fish every 12 hours, in order to confirm complete anesthetization during development. In the rare cases motion is observed in the tricained fish (~4 hours on Day 3 or ~ 9 hours on Day 5), we also observed motion in the wells in which we did not place fish, indicating that we are experiencing either shot noise or motion in the rig room, rather than observing tricainated fish that have "woken up".

**Supplemental Movie 1**

We illustrate the on-rate of tricaine by applying anesthesia to a fish reared under standard conditions. We pour the anesthesia on at 10s, and for the following 20s the fish slows its swim bouts. After 30-40s, the fish floats, losing posture and no longer responding to physical stimuli.

**Supplemental Movie 2**

We illustrate the off-rate of tricaine by washing out the anesthesia in the fish in Supplemental Movie 1. In the first 10s, the fish is in a tricaine bath, and continues to be immobile and does not respond to physical touch. We then transfer the fish to a bath of standard fish water by 20s, and after 1 minute and 10 seconds we see the first touch-evoked swim. The fish continues to recover for the next minute, and by 2 minutes after wash out begins to swim in responses to taps.

**Supplemental Movie 3**

Recording of fish in Figure 4 while still under tricaine (first 303 frames, 10s at 30 frames/s) and after washout (frames 304-1201, last 30s at 30 frames/s). No coordinated activity is observed while still under anesthesia, despite being shown continuous visual stimuli, while strong coordinated activity is seen even immediately after washout.

**Supplemental Videos are also available at**
**https://www.dropbox.com/sh/dgleogp77q8z2dz/AACyDixGe5TPV4JnBYbSwcBPa?dl=0**
